# An improved ATAC-seq protocol reduces background and enables interrogation of frozen tissues

**DOI:** 10.1101/181206

**Authors:** M. Ryan Corces, Alexandro E. Trevino, Emily G. Hamilton, Peyton G. Greenside, Nicholas A. Sinnott-Armstrong, Sam Vesuna, Ansuman T. Satpathy, Adam J. Rubin, Kathleen S. Montine, Beijing Wu, Arwa Kathiria, Seung Woo Cho, Maxwell R. Mumbach, Ava C. Carter, Maya Kasowski, Lisa A. Orloff, Viviana I. Risca, Anshul Kundaje, Paul A. Khavari, Thomas J. Montine, William J. Greenleaf, Howard Y. Chang

## Abstract

We present Omni-ATAC, an improved ATAC-seq protocol for chromatin accessibility profiling that works across multiple applications with substantial improvement of signal-to-background ratio and information content. The Omni-ATAC protocol enables chromatin accessibility profiling from archival frozen tissue samples and 50 μm sections, revealing the activities of disease-associated DNA elements in distinct human brain structures. The Omni-ATAC protocol enables the interrogation of personal regulomes in tissue context and translational studies.

## MAIN TEXT

The mapping of regulatory landscapes that control gene expression and cell state has become a widespread area of interest. Recent methodological advances such as the advent of the assay for transposase-accessible chromatin by sequencing^1^ (ATAC-seq) and the application of DNase hypersensitivity sequencing (DNase-seq) to low cell numbers^2^ have enabled the generation of high-fidelity chromatin accessibility profiles for a variety of cell types^3–9^. However, certain cell types and tissues require individualized protocol optimizations^10,11^, making data difficult to compare across multiple studies. To this end, we report an improved, broadly applicable ATAC-seq protocol, called Omni-ATAC, which is suitable for diverse cell lines, tissue types, and archival frozen samples while simultaneously improving data quality across all cell types and cell contexts tested (**Supplementary Protocol 1**).

Systematic protocol alterations lead to stepwise improvements in ATAC-seq data quality while maintaining the simplicity of the standard ATAC-seq protocol (Supplementary Fig. 1a). These improvements include (i) the use of multiple detergents (NP40, Tween-20, and digitonin) to improve permeabilization across a wide array of cell types and remove mitochondria from the transposition reaction, (ii) a post-lysis wash step using Tween-20 to further remove mitochondria and to increase the library complexity, and (iii) the use of PBS in the transposition reaction to increase the signal-to-background ratio (**Supplementary** Fig. 1a and **Supplementary Note**). ATAC-seq data generated using the Omni-ATAC protocol are consistent with previously published standard ATAC-seq^1^ (R=0.73), Fast-ATAC^11^ (R=0.88), and DNase-seq^12^ (R=0.72) measurements in GM12878 and CD4+ T cells (**Supplementary** Fig. 1b-i and **Supplementary** Fig. 2a-f). However, compared to the standard ATAC-seq protocol, the Omni-ATAC protocol lowers sequencing costs by generating 13-fold fewer sequencing reads mapping to mitochondrial DNA, and improves data quality by yielding 3-fold higher percentage of reads mapping to peaks of chromatin accessibility and 15-fold greater number of unique fragments per input cell (median values from N=14 cell types/contexts; all values determined from 5 million random aligned de-duplicated reads; Fig. 1a, **Supplementary Table 1**, and **Supplementary Fig. 3a**). Of the sequencing reads mapping to known peaks, the Omni-ATAC protocol generates a higher percentage of both reads mapping to promoters, defined as within 500 bp of a transcriptional start site (TSS), and reads mapping to distal elements, defined as more than 500 bp away from a TSS, as compared to the standard ATAC-seq, Fast-ATAC, and DNase-seq methodologies (Fig. 1b and **Supplementary Fig. 3b**). With more information per sequencing read, the Omni-ATAC protocol identifies as many or more peaks with consistently higher significance at constant sequencing depth than previously published standard ATAC-seq, Fast-ATAC, and DNase-seq methodologies (**Supplementary Fig. 3c-e**). Of the peaks identified by at least two methods, 53.8% are identified by all methods, 38.5% are not identified by the standard ATAC-seq method, 6.0% are not identified by DNase-seq, and 1.7% are not identified by the Omni-ATAC protocol (Fig. 1c).This demonstrates that both the Omni-ATAC protocol and DNase-seq identify a substantial number of peaks that are neither identified, nor show robust signal, using the standard ATAC-seq protocol (**Supplementary Fig. 3f-h**). Stronger signals can clearly be observed in sequencing tracks derived from equivalent numbers of non-duplicate, aligned reads (**Supplementary Fig. 3i-l**), demonstrating that the Omni-ATAC protocol produces accessibility data with substantially higher signal-to-background ratio than alternative tested methods.

**Figure 1.**
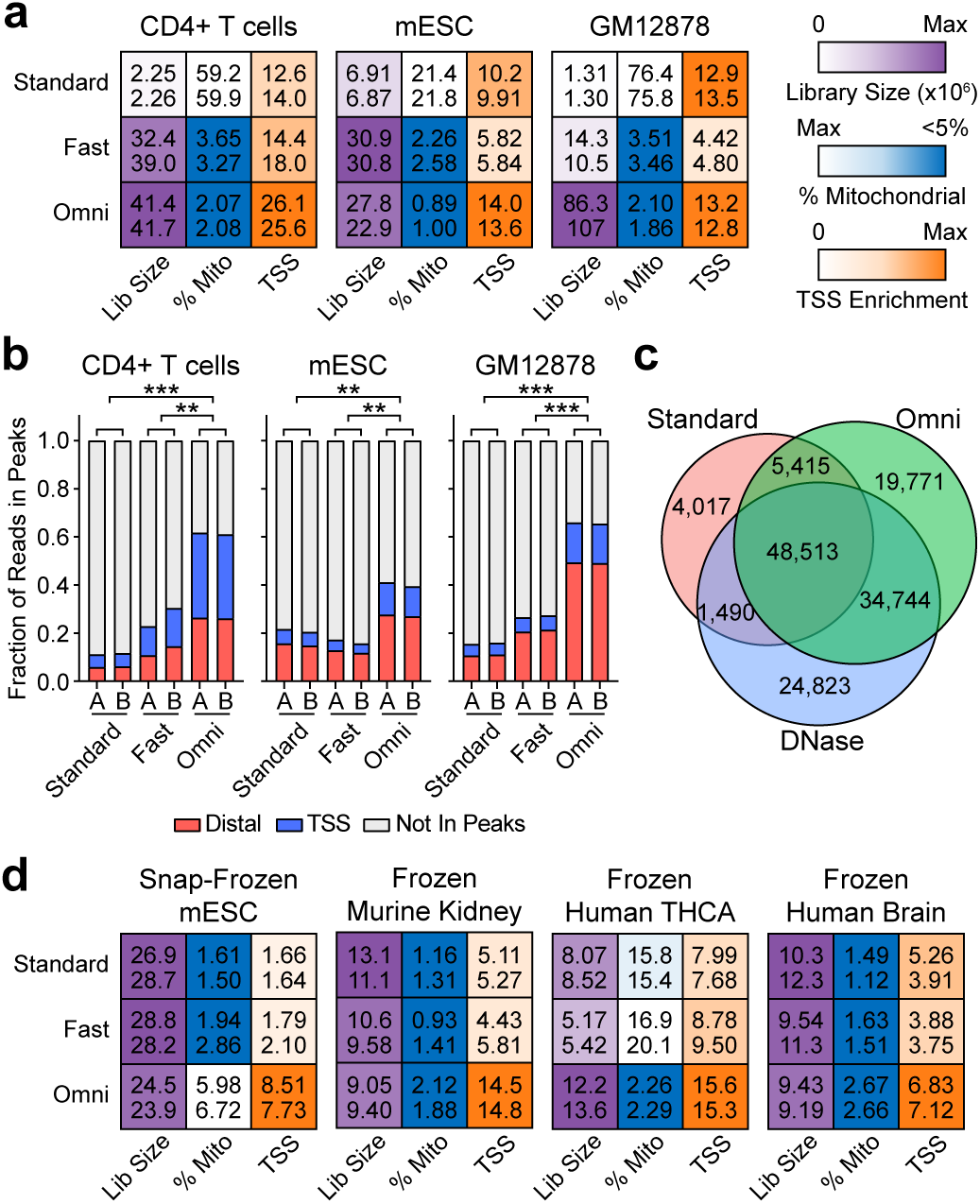
Omni-ATAC protocol compared to standard ATAC-seq or Fast-ATAC. (**a**) Heatmap-based representation of ATAC-seq quality control metrics including library size (purple), percent of reads mapping to mitochondrial DNA (blue), and enrichment of signal at TSSs (orange). Library size (“Lib Size”), percent of reads mapping to mitochondrial DNA (“% Mito”), and the TSS enrichment score (“TSS”) are shown. Deeper color is used to depict the most desirable value of each statistic and ranges following a linear scale starting at 0 (white) and ending at the maximum value for the given cell type. In the case of the percent of reads mapping to chrM, the color scale starts at the maximum value (white) and ends at 5% (blue). All values were determined from 5 million random aligned reads. Two technical replicates are shown per sample as numeric values within each box. (**b**) Fraction of reads in peaks mapping to TSSs (+/- 500 bp of TSS) and distal elements (> 500 bp from TSS) from libraries generated in this study using the standard ATAC-seq method (“Standard”), the Fast-ATAC-seq method (“Fast”) or the Omni-ATAC protocol (“Omni”). Each bar represents a single technical replicates. **p< 0.01; ***p< 0.001 by two-tailed unpaired students t-test comparing the fraction of reads in peaks to reads outside of peaks. All values were determined from 5 million random aligned de-duplicated reads. (**c**) Overlap of GM12878 peaks called from 60 million reads using standard ATAC-seq, DNase-seq, and Omni-ATAC. All input reads were trimmed to equal length (36 bp) prior to alignment. Data represents the mean of three individual down-sampling replicates. (**d**) Heatmap-based representation of ATAC-seq quality control metrics including library size, percent of reads mapping to mitochondrial DNA, and enrichment of signal at TSSs as shown in (a). Deeper color is used to depict the most desirable value of each statistic. All values were determined from 5 million random aligned reads. Two technical replicates are shown per sample as numeric values within each box.

**Figure 2.**
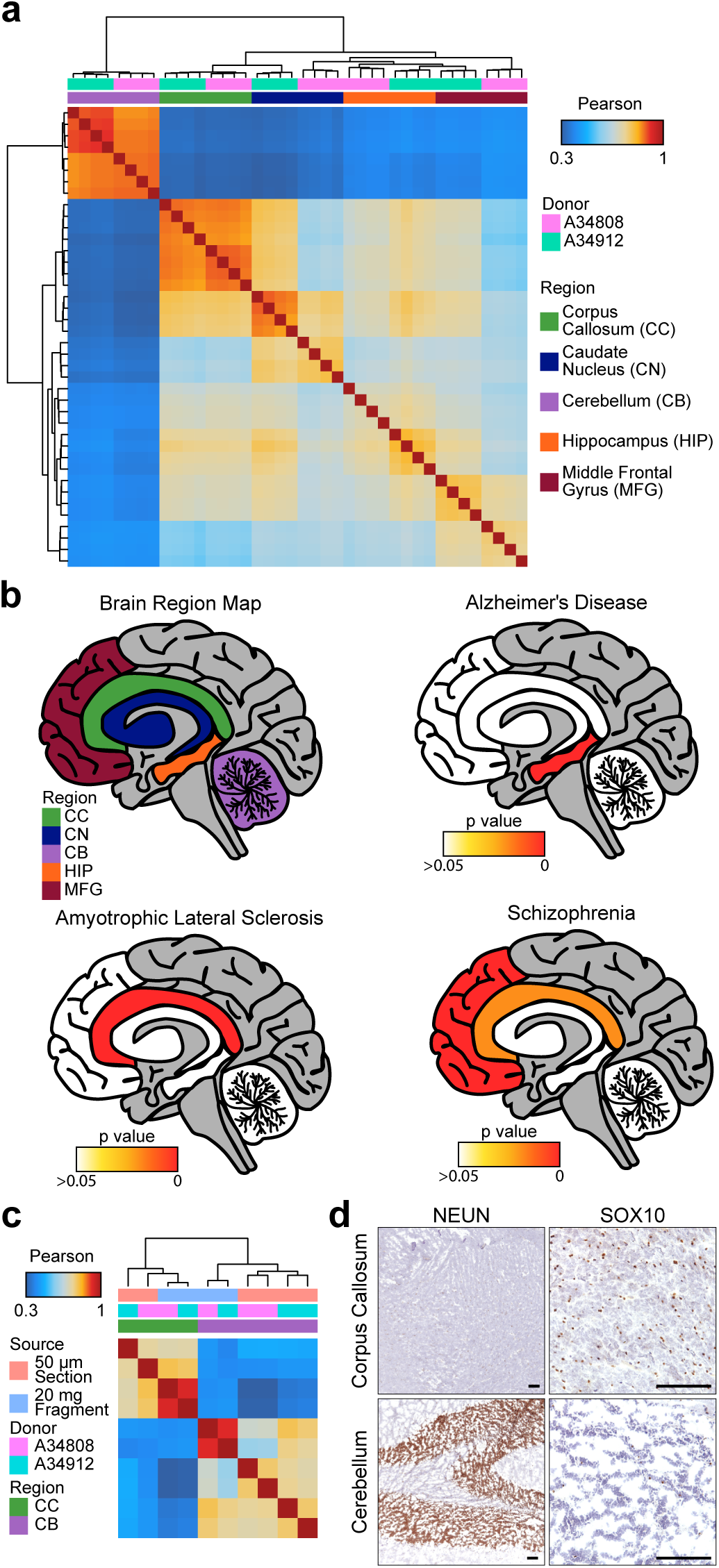
Omni-ATAC in defined post-mortem human brain. (**a**) Pearson correlation heatmap showing sample by sample unsupervised clustering on all peaks identified across all regions. Biological donor and brain region are indicated by color across the top. CC = corpus callosum; CN = caudate nucleus; CB = cerebellum; HIP = hippocampus; MFG = middle frontal gyrus. All regions are represented by four technical replicates per donor. (**b**) Significance of enrichment of disease-specific GWAS polymorphisms in the uniquely-accessible regions of the 5 different brain regions shown at the top left. The empirical p value is depicted colorimetrically with reference to association-based permutations of the GWAS SNPs. Non-significant empirical p values are represented by white coloration. (**c**) Pearson correlation heatmap showing sample by sample unsupervised clustering on all peaks. Data derived from 20 mg tissue fragments represents the mean of 4 technical replicates from a single biological donor. Data derived from 50 μm tissue sections shows individual technical replicates. (**d**) Histology and immunohistochemistry of 5 μm frozen sections from cerebellum and corpus callosum immediately adjacent to the 50 μm section used for ATAC-seq shown in Supplementary Figure 8. Stains from left to right: anti-NEUN immunohistochemistry, and anti-SOX10 immunohistochemistry. Scale bar represents 100 μm in each image. SOX10 staining shown at higher resolution due to more diffuse staining pattern.

The Omni-ATAC protocol also improves chromatin accessibility measurements from very small numbers of cells. Previous publications have demonstrated the applicability of standard ATAC-seq to as few as 500 cells^1,6^. ATAC-seq using the Omni-ATAC protocol in 500 GM12878 cells leads to a significant increase in signal-to-background ratio and fraction of reads in peaks compared to previously published data^1^ (**Supplementary Fig. 4a-g,** and **Supplementary Table 1**). Moreover, ATAC-seq using the Omni-ATAC protocol in 500 GM12878 cells identifies more known accessible chromatin regions (**Supplementary Fig. 4h**) and shows a greater correlation with libraries generated from 50,000 cells than standard ATAC-seq (**Supplementary Fig. 4i,j**). We also note that the Omni-ATAC protocol has the potential to reduce reaction costs by enabling the use of less transposase enzyme (**Supplementary Note, Supplementary Figure 5a-e,** and **Supplementary Table 1**).

In addition to improving data quality in previously studied cell types, the Omni-ATAC protocol enables the generation of robust chromatin accessibility data from cell types and cell contexts that previously proved difficult to assay. For example, standard ATAC-seq and Fast-ATAC generally perform poorly on snap-frozen pellets, requiring the transposition reaction to be performed on fresh cells. This constraint has prevented the broad application of ATAC-seq to banked frozen pellets. However, the Omni-ATAC protocol allows for the generation of high signal-to-background chromatin accessibility profiles from snap-frozen pellets of just 50,000 cells (Fig. 1d and **Supplementary Table 1**). Similarly, although both standard ATAC-seq and Fast-ATAC perform poorly on primary human keratinocytes, yielding data with low signal-to-background and a low fraction of reads in peaks, the Omni-ATAC protocol allows for the generation of high-quality, information-rich chromatin accessibility data under a single consistent protocol (**Supplementary Table 1**). Overall, the Omni-ATAC protocol simplifies laboratory workflows and enables data acquisition from biomaterials previously deemed unusable for native chromatin accessibility profiling.

We sought to generate high-quality chromatin accessibility profiles from clinically relevant frozen tissues such as brain, where clinical specimens are acquired from rapid autopsy and preserved by snap-freezing, or cancer, where patients’ tissue samples are often banked as snap-frozen fragments. We first isolated nuclei from 20 mg of frozen tissue via Dounce homogenization followed by density gradient centrifugation (**Supplementary Protocol 2** and **Supplementary Fig. 6a,b**). The Omni-ATAC protocol provided improved signal-to-background and overall data quality in frozen human thyroid cancer (THCA), frozen post-mortem human brain samples, and a diverse array of frozen mouse tissues including colon, heart, liver, lung, spleen, and kidney (Fig. 1d and **Supplementary Table 1**). We then applied this method to study diverse macro-dissected human brain regions including the cerebellum, caudate nucleus, corpus callosum, middle frontal gyrus, and hippocampus from two donors (**Supplementary Table 2**). Comparison of the 5 brain regions (**Supplementary Table 1**) shows strong intra-region correlation across technical replicates and biological donors (Fig. 2a and **Supplementary Fig. 6c**), allowing delineation of region-specific signatures of differentially accessible chromatin and differential transcription factor motif usage that correlate with known brain-specific transcriptional drivers^13^ (**Supplementary Fig. 6d,e**).

These chromatin accessibility profiles also enable interpretation of the results of genome-wide association studies (GWAS) that have mapped putative brain disease-relevant SNPs to non-coding regions (Fig. 2b, **Supplementary Fig. 7a-c**, and **Supplementary Table 3**). For example, the hippocampus, a region that plays key roles in memory formation and exhibits atrophy in Alzheimer’s disease^14^, shows the strongest enrichment for Alzheimer’s disease-related GWAS SNPs (Fig. 2b and **Supplementary Table 3**). Similarly, the corpus callosum, a region that is consistently involved in amyotrophic lateral sclerosis (ALS)^15^, shows significant enrichment for ALS-related GWAS SNPs^16^ (Fig. 2b, **Supplementary Fig. 8a** and **Supplementary Table 3**).

Given the potential applicability of epigenomic information to clinical diagnosis and prognostication, we developed a methodological framework to combine routine histo-pathology with sub-millimeter precision ATAC-seq. This approach enables collection of multiple thin 5 μm tissue sections for pathology immediately adjacent to a single 50 μm tissue section used for ATAC-seq. On 50 μm frozen tissue sections from human brain regions, the Omni-ATAC protocol generates chromatin accessibility profiles comparable to those generated from bulk tissue (Fig. 2c, **Supplementary Fig. 8b-f**, and **Supplementary Table 1**). These chromatin accessibility profiles correlate well with adjacent histopathological staining with regions of high glial cell abundance by SOX10 immunohistochemistry showing increased accessibility near glial-specific genes such as *OLIG2* (Fig. 2d, **Supplementary Fig. 8b,c,g** and **Supplementary Fig. 9a-d**) and regions of high neuronal cell abundance by NEUN immunohistochemistry and Nissl staining showing increased accessibility near neuron-specific genes such as *NEUROD1* (Fig. 2d, **Supplementary Fig. 8d,g,** and **Supplementary Fig. 9e-h**). Thus, the Omni-ATAC protocol enables the application of epigenomics to clinically relevant specimens, paving the way for assays and diagnostics that leverage the highly informative and cell type-specific signals of the open chromatin landscape.

Altogether, our data demonstrate that the Omni-ATAC protocol provides a robust, broadly applicable platform for the generation of high-quality and information-rich chromatin accessibility data. By enabling profiling in a wider array of cell types and cell contexts, we believe the Omni-ATAC protocol will make chromatin accessibility landscapes more universally comparable and facilitate the use of ATAC-seq in difficult cell lines, rare primary cells, and otherwise complicated applications such as snap-frozen cell pellets and clinically relevant tissues.

## DATA AVAILABILITY

All raw sequencing files generated in this work are available through the Sequencing Read Archive via BioProject PRJNA380283 or SRA SRP103230.

## ACKNOWLEDGEMENTS

We thank the Stanford Alzheimer’s Disease Research Center (NIH P50 AG047366), the Pacific Udall Center for Excellence in Parkinson’s Disease Research (NIH P50 NS062684) and their participants for donating samples for research. We thank John Coller and Xuhuai Ji for sequencing assistance. We thank Ed Plowey and Divya Channappa for tissue preparation, and Pauline Chu and Amarjeet Grewall for histology assistance. M.R.C. is supported by a grant from The Leukemia & Lymphoma Society Career Development Program and NIH training grant R25CA180993. A.E.T. is supported by the National Science Foundation Graduate Research Fellowship Program. This research was conducted with Government support under and awarded by DoD, Air Force Office of Scientific Research, National Defense Science and Engineering Graduate (NDSEG) Fellowship, 32 CFR 168a to N.A.S.A. Supported by NIH P50-HG007735 (to H.Y.C. and W.J.G.) and NIA RF1 AG053959 (to T.J.M.).

## AUTHOR CONTRIBUTIONS

M.R.C. and H.Y.C. conceived the project. M.R.C., E.G.H., and A.T.S. performed all experiments with help from M.R.M., and A.C.C. S.W.C produced the Tn5 transposase complex used in all experiments. P.G.G. performed all GWAS analysis with guidance and supervision from A.K. M.R.C. and A.J.R performed all other data analysis. M.R.C., A.E.T., S.V., and N.A.S-A., developed methods for the isolation of nuclei from frozen tissues. B.W., A.K., V.I.R., and W.J.G provided protocol expertise and recommendations. K.S.M. and T.J.M. oversaw all brain tissue acquisition and processing. M.K. and L.A.O. oversaw acquisition and processing of thyroid cancer tissue. A.J.R. and P.A.K. oversaw acquisition and processing of primary foreskin keratinocytes. M.R.C., E.G.H., W.J.G., and H.Y.C. wrote the manuscript with input from all authors.

## COMPETING FINANCIAL INTERESTS

H.Y.C. and W.J.G. are co-founders and consultants of Epinomics. A.K. is a member of the scientific advisory board of Epinomics.

## ONLINE METHODS

### Code Availability

All custom code used in this work is available upon request.

### Publicly Available Data Used In This Work

GM12878 standard ATAC-seq data was obtained as raw fastq files from GEO GSE47753. GM12878 DNase-seq data was obtained as unfiltered and filtered alignments from ENCODE ENCSR000EMT. mESC ES-14 DNase-seq data was obtained as filtered alignments from ENCODE ENCSR000CMW.

### Genome Annotations

All human data is aligned and annotated for the hg19 reference genome. All murine data is aligned and annotated for the mm10 reference genome.

### Sequencing

All deep sequencing was performed using 2x75 bp reads on an Illumina HiSeq4000 instrument that was purchased with funds from the NIH under award number S10OD018220. Prior to sequencing on a HiSeq4000 pooled ATAC-seq libraries were purified using PAGE gel size selection (for fragments > 100 bp) to remove excess primers (< 100 bp). All low depth sequencing data was performed using 2x75 bp reads on an Illumina MiSeq instrument.

### Sample acquisition and patient consent

Primary blood cells, primary brain tissue, primary thyroid cancer tissue, and primary keratinocytes were acquired with written and informed consent through Stanford IRB protocols 27804, 29259, 11977, and 35324 respectively. Human donor sample sizes were chosen to provide sufficient confidence to validate methodological conclusions of the applicability of Omni-ATAC. All animal studies were performed in compliance with IACUC and APLAC regulations.

### Omni-ATAC protocol

See Supplementary Protocol 1 and 2 for a detailed protocol. Cells grown in tissue culture were pre-treated with 200 U/ml DNase (Worthington) for 30 minutes at 37°C to remove free-floating DNA and digest DNA from dead cells. This media was then washed out and the cells were resuspended in cold PBS. For primary human T cells, cells were sorted using a Becton Dickinson FACS Aria II based on the expression of CD45, CD3, and CD4 as described previously^11^. After counting, 50,000 cells were resuspended in 1 ml of cold ATAC-seq Resuspension Buffer (RSB - 10 mM Tris-HCl pH 7.4, 10 mM NaCl, 3mM MgCl2 in water). Cells were centrifuged at 500 RCF for 5 minutes in a pre-chilled 4°C fixed-angle centrifuge. After centrifugation, 900 ul of supernatant was aspirated, leaving 100 ul of supernatant. This remaining 100 ul of supernatant was carefully aspirated by pipetting with a p200 pipette, avoiding the cell pellet. Cell pellets were then resuspended in 50 ul of ATAC-seq RSB containing 0.1% NP40, 0.1% Tween-20, and 0.01% Digitonin by pipetting up and down 3 times. This cell lysis reaction was incubated on ice for 3 minutes. After lysis, 1 ml of ATAC-seq RSB containing 0.1% Tween-20 (without NP40 or digitonin) was added and tubes were inverted to mix. Nuclei were then centrifuged for 10 minutes at 500 RCF in a pre-chilled 4°C fixed-angle centrifuge. Supernatant was removed as before with two pipetting steps and nuclei were resuspended in 50 ul of transposition mix (25 ul 2x TD buffer, 2.5 ul transposase (100nM final), 16.5 ul PBS, 0.5 ul 1% digitonin, 0.5 ul 10% Tween-20, 5 ul H2O) by pipetting up and down 6 times. Transposition reactions were incubated at 37°C for 30 minutes in a thermomixer with 1000 RPM shaking. Reactions were cleaned up with Zymo DNA Clean and Concentrator 5 columns. The remainder of the ATAC-seq library preparation was performed as described previously^17^. All libraries were amplified with a target concentration of 20 ul at 4 nM which is equivalent to 80 femtomoles of product. Minor protocol modifications were used for Omni-ATAC in frozen tissues and in limiting cell numbers. These modifications are outlined in the corresponding methods sections.

### Other ATAC-seq Methodologies

Standard ATAC-seq and Fast-ATAC were performed as described previously^1,11^ without additional modifications.

### ATAC-seq of frozen cell pellets with the Omni-ATAC protocol

In this work, frozen cell pellets of 50,000 cells were directly transposed using the Omni-ATAC transposition mix without additional lysis and wash steps. However, depending on the cell type and application, it may improve data quality to thaw the cell pellet in Omni-ATAC lysis buffer, wash, and then transpose.

### 500 cell ATAC-seq with the Omni-ATAC protocol

GM12878 cells were counted five times using a manual hemocytometer. The mean cell count was used to resuspend the cells to a concentration of 500 cells per 100 ul by addition of PBS. From this diluted cell mixture 100 ul (500 cells) were deposited into a 0.5 ml DNA LoBind tube (Eppendorf #022431005) containing 400 ul of cold ATAC-seq RSB. This was done to simulate a workflow involving FACS sorting. These tubes were centrifuged at 500 RCF for 10 minutes in a pre-chilled 4°C fixed-angle centrifuge with 0.6 ml tube adapters. All supernatant was removed using two pipetting steps as described above, first removing 400 ul with a p1000 pipette followed by removal of the remaining volume with a p200 pipette. We note that a gradual but constant removal of supernatant is crucial and the final supernatant removal should be completed in a single motion to avoid disrupting the cell pellet. After supernatant removal, lysis and transposition were performed simultaneously to avoid cell loss and the total reaction volume was reduced for the same reason. As such, 10 ul of transposition mix (3.3 ul PBS, 1.15 ul H2O, 5 ul 2x TD Buffer, 0.25 ul 1:10 diluted Tn5, 0.1 ul 1% Digitonin, 0.1 ul 10% Tween-20, 0.1 ul 10% NP40) was added directly to the not-visible cell pellet and the pellet was resuspended by pipetting up and down 6 times. The transposition reaction was incubated at 37°C for 30 minutes in a thermomixer with 1000 RPM shaking. Tn5 should be diluted in 1x TD Buffer (for example, 5 ul 2x TD Buffer, 4 ul of water, 1 ul Tn5).

### ATAC-seq using the Omni-ATAC protocol on Nuclei Isolated from Frozen Tissues

See Supplemental Protocol 2 for a detailed protocol. This protocol is highly similar to the INTACT method^18^ and either can be used for nuclei isolation with equivalent results. All steps were carried out at 4°C. A frozen tissue fragment approximately 20 mg was placed into a pre-chilled 2 ml Dounce homogenizer containing 2 ml of cold 1x homogenization buffer (320 mM sucrose, 0.1 mM EDTA, 0.1% NP40, 5mM CaCl2, 3mM Mg(Ac)2, 10 mM Tris pH 7.8, 1x protease inhibitors (Roche, cOmplete), 167 uM β-mercaptoethanol, in water). Tissue was homogenized with approximately 10 strokes with the loose “A” pestle, followed by 20 strokes with the tight “B” pestle. Connective tissue and residual debris were pre-cleared by filtration through an 80 μm nylon mesh filter followed by centrifugation for 1 min at 100 RCF. Avoiding pelleted debris, 400 ul was transferred to a pre-chilled 2 ml round bottom Lo-Bind Eppendorf tube. An equal volume (400 ul) of a 50% iodixanol solution (50% iodixanol in 1x homogenization buffer) was added and mixed by pipetting to make a final concentration of 25% iodixanol. 600 ul of a 29% iodixanol solution (29% iodixanol in 1x homogenization buffer containing 480 mM sucrose) was layered underneath the 25% iodixanol mixture. A clearly defined interface should be visible. In a similar fashion, 600 ul of a 35% iodixanol solution (35% iodixanol in 1x homogenization containing 480 mM surcose) was layered underneath the 29% iodixanol solution. Again, a clearly defined interface should be visible between all three layers. In a swinging bucket centrifuge, nuclei were centrifuged for 20 minute at 3000 RCF. After centrifugation, nuclei are present at the interface of the 29% and 35% iodixanol solutions. This nuclei band was collected in a 300 ul volume and transferred to a pre-chilled tube. Nuclei were counted after addition of trypan blue which stains all nuclei due to membrane permeabilization from freezing. 50,000 counted nuclei were then transferred to a tube containing 1 ml of ATAC-seq RSB with 0.1% Tween-20. Nuclei were pelleted by centrifugation at 500 RCF for 10 minutes in a pre-chilled 4°C fixed-angle centrifuge. Supernatant was removed using two pipetting steps as described above. As nuclei are already permeabilized, no lysis step was performed and the transposition mix (25 ul 2x TD buffer, 2.5 ul transposase (100nM final), 16.5 ul PBS, 0.5 ul 1% digitonin, 0.5 ul 10% Tween-20, 5 ul H2O) was added directly to the nuclear pellet and mixed by pipetting up and down 6 times. Transposition reactions were incubated at 37°C for 30 minutes in a thermomixer with 1000 RPM shaking. Reactions were cleaned up with Zymo DNA Clean and Concentrator 5 columns. The remainder of the ATAC-seq library preparation was performed as described previously^17^.

### ATAC-seq using the Omni-ATAC protocol on Thin Frozen Tissue Sections

Omni-ATAC on thin frozen tissue sections was performed using the same protocol as for 20 mg tissue fragments with one modification. To prevent sample loss, 50 μm sections were prepared in a 2 ml Dounce homogenizer containing 500 ul of 1x homogenization buffer. We determined that, despite some bubble formation, the quality of nuclei recovered from homogenization in a 2 ml Dounce with 500 ul is superior to the quality of nuclei recovered when smaller Dounce homogenizers were used.

### ATAC-seq Data Analysis

ATAC-seq data analysis used the following tools and versions: Samtools v1.3, Picard v2.2.1, Bowtie2 v2.2.8, macs2 v2.1.0.20150731, bedtools v2.23.0. First, Nextera adapter sequences were trimmed from reads using a custom python script. These reads were aligned to a reference genome using bowtie2 with standard parameters and a maximum fragment length of 2000. Picard was then used to remove duplicate reads. These de-duplicated reads were then filtered for high quality (MAPQ > = 30), non-ChrM/ChrY, and properly paired (samtools flag 0x2) reads.

### ATAC-seq Library QC Statistics

Library size is determined from a subsample of 5 million aligned reads prior to any filtration. Subsampling to 5 million reads was used because current library size estimate tools are very sensitive to the input read depth. In this way, because the library size estimates are obtained from the same number of input reads, our library size estimates are comparable across assays and cell types but may be an underestimate of the actual library complexity as only 5 million reads were used. The percent of reads aligning to mitochondrial DNA and the transcriptional start site enrichment were also determined from the same 5 million read subset or aligned reads. Transcription start site enrichment was determined using hg19 RefSeq TSSs. Enrichment was calculated by counting transposition events in 1 base pair bins in the regions +/- 2000 bp surrounding all TSSs. The value of each bin was then normalized by dividing by the mean value of the first 200 single base pair bins. In this way, the signal at bases −2000:-1800 is used to represent the “background” signal. For low depth libraries presented in Supplementary Table 1, only 100,000 aligned reads were used for these metrics.

### Footprinting Analysis

Meta-footprints were generated for CTCF using pyDNase, a tool based on the Wellington algorithm^19^. CTCF motif occurrences were filtered for those sites overlapping an ATAC-seq peak with a peak score (-log10(pvalue)) greater than 50. The resulting high-confidence CTCF motif set was used as input to pyDNase dnase_average_profile.py using the –c (stranded) and –A (accounts for Tn5 cut-site offset) options. Meta-footprints were generated from all available filtered reads.

### Fraction of Reads in Peaks (FRIP)

The fraction of reads in peaks was determined using a subsample of 5 million aligned de-duplicated reads prior to any filtration. For FRIP calculations, called peaks were marked as “distal” if they showed no overlap with +/- 500 bp from annotated transcriptional start sites. Reads overlapping with regions +/- 500 bp from TSSs were binned as “TSS” reads. Reads overlapping with distal peaks were binned as “distal” reads. All other reads not overlapping one of these regions were labeled as “Not in peaks”. Overlap of reads with genomic regions was determined using bedtools intersect with standard parameters. Reads mapping to mitochondrial DNA are categorically binned as “Not in peaks”. For low depth libraries presented in Supplementary Table 1, only 100,000 aligned reads were used for these metrics. Peak files used for FRIP calculations are outlined in Supplementary Table 1.

### Peak Calling and Peak Scores

All peak calling was performed with macs2 using “macs2 callpeak --nomodel --nolambda --keep-dup all --call-summits”. For simulations of peaks called per input read, aligned and de-duplicated BAM files were used without any additional filtering.

### Peak Overlap and Venn Diagrams

For peak overlap of DNase, standard ATAC-seq, and Omni-ATAC-seq, peaks were called using fully processed filtered and merged BAM files representing the union of all available replicates. These individual peak sets were concatenated and a union peak set was made as described previously^11^. Briefly, overlapping peaks called in different data sets were resolved by retaining the peak with the higher macs2 score. In this way, we generated a non-overlapping union peak set containing all of the peaks called in data from all three assays. This union peak set was then intersected individually with each of the peak sets for DNase/standard ATAC-seq/Omni-ATAC-seq. Each individual intersection represented the total peaks called in each individual assay. These peak sets were then intersected with each other to determine the overlap of peaks called. All intersections were performed using bedtools^20^ with either the “-v” (unique) and the “-u” (shared) options.

### Sequencing Tracks

All sequencing tracks were made using the WashU Epigenome Browser. Sequencing coverage tracks used to compare DNase, standard ATAC-seq, and Omni-ATAC-seq were generated by subsampling 60 million reads from an aligned and de-duplicated BAM file that has not been additionally filtered. These equal-depth BAM files were then converted to bigwig for visualization. For comparisons involving DNase-seq, all ATAC-seq reads were trimmed to 36 bp to match the single-end 36 bp sequencing used in DNase prior to alignment. The y-axis scale for all sequencing tracks was set to range from 0 to the maximum height among the three datasets. In this way, the heights of the tracks are comparable across techniques as they are derived from the same number of equal-length input reads. Sequencing tracks used to compare 500 cell Omni-ATAC to 500 cell standard ATAC-seq in GM12878 cells were not normalized. In these visualizations, all pass filter reads were used to generate sequencing tracks under the assumption that these libraries were sequenced to near full depth. This is due to differences in the library sizes of the Omni-ATAC and standard ATAC-seq 500 cell libraries. Sequencing tracks related to frozen human brain tissue are all normalized by the total number of reads in peaks.

### GWAS Enrichment

To test the for enrichment of GWAS variants in our region-specific uniquely accessible peak sets (Supplementary Fig. 6d), we used all GWAS data sets in the GRASP database (n=178)^21^. The GWAS SNPs were pruned to contain no variants in linkage disequilibrium by keeping the most significant p-value where there were multiple linked variants for the same trait. We only kept GWAS with at least 900 SNPs after pruning in the analysis for sufficient quality to calculate an enrichment. The set of pruned SNPs was then expanded to all linked variants with European r^2^ ≥ 0.8 for all further analysis.

We performed a rank-based enrichment of GWAS variants in each set of up-regulated differential peaks and merged broad and narrow peaks of each brain region. We segmented each GWAS study into bins representing decreasing tiers of significance. We set a minimum bin size of 50 and filled the first bin with the 50 most significantly associated variants for each study. We then filled the next bins with 2*50, 4*50 and 8*50 variants and then segmented the remaining variants into bins at the four quartiles of the remaining p-value distribution. We used the pruned set of SNPs for setting the bin thresholds. We then computed the rank fold change enrichment of ATAC peaks across the segmented GWAS^22^. For each bin we computed the fraction of GWAS variants less than or equal to the bin’s p-value threshold that overlapped the peaks for each brain region. We calculated the fold change enrichment by dividing this fraction by the fraction of all GWAS variants of any significance level overlapping our regions. Baseline enrichment is 1 indicating no change from the base rate of overlap of all significant and non-significant variants in the study. An enrichment less than 1 means the most significant variants are depleted relative to the baseline and any value greater than 1 indicates significant variants are enriched. To compute the significance of these enrichments, we permuted the p-value associated with each GWAS SNP in the study 200 times and re-computed the enrichment relative to baseline. The empirical p-value indicates the number of permuted studies where the true study has a greater enrichment for the most significant bin of GWAS hits.

### Correlation Analyses

Pearson correlation heatmaps were generated using variance stabilizing transformed read counts via DESeq2^23^. Briefly, read counts were determined for all called peaks using bedtools multicov and then quantile normalized and rounded to the nearest read count. This normalized count matrix was input to DESeq2. Plots of sample by sample correlation at all peaks were generated based on quantile normalized read counts.

### ATAC-seq Peak and TF Analyses

Brain region-unique peaks were identified using quantile normalized read counts. Briefly, peaks were identified as unique if they were found to be more than two standard deviations away from the mean of all other brain regions. Transcription factor deviation analysis was performed as described previously^24^. Enrichment of TF motifs in ATAC-seq peak regions was also performed using the Hypergeometric Optimization of Motif Enrichment (Homer) algorithm with standard parameters.

### Histology and Immunohistochemistry

Histology and immunohistochemistry were performed according to manufacturer protocols. Anti-NEUN (ab177487) and anti-SOX10 (ab212843) antibodies were purchased from Abcam.

### Transposase Production

All transposase used in this work was produced and prepared as described previously^25^.

### Cell Lines

All cell lines used in this study were purchased from ATCC or DSMZ. Where possible, cell lines were validated by comparison to published sequencing data or by in-house genotyping with comparison to the Cancer Cell Line Encyclopedia. Cell lines were tested for mycoplasma contamination upon receipt and periodically thereafter but not prior to each experiment.

### Statistics

All statistical tests performed are included in the figure legends where relevant.

